# Digital Twins for Fungal Computing: Viable XOR Regimes, Parameter Inference, and Waveform-Guided Rediscovery

**DOI:** 10.64898/2026.03.27.714860

**Authors:** Kiran Bhattacharyya

**Author notes:** Data/Code available at http://your.repo.here.com.

## Abstract

Fungal substrates are promising candidates for unconventional computing, but specimen-to-specimen variability makes logic-gate fabrication difficult to reproduce. This paper presents a digital-twin workflow for fungal excitable networks and evaluates three components needed for computeraided design: identifying parameter regimes that support XOR computation, inferring latent biophysical parameters from electrical characterization data, and refining those inferred parameters by waveform matching. The model represents mycelium as a random geometric graph with FitzHugh–Nagumo node dynamics and memristive edge conductances. A systematic optimization study over 160 simulated specimens identifies a viable XOR subspace defined by tuned biophysical parameters, electrode geometry, and stimulus timing. A characterization study over 400 simulated specimens uses step-response, paired-pulse, and triangle-sweep protocols to extract 94 response features. Random forest regressors recover several latent parameters reliably (*R*^2^ = 0.912 for *τ*_*v*_, 0.816 for *τ*_*w*_, 0.717 for *a*), while *v*_scale_, *R*_on_, and *R*_off_ remain weakly identifiable. On a preliminary rediscovery validation using 15 optimized specimens (20–50 nodes), ML initialization followed by local waveform-matching refinement reduces mean waveform mismatch from 1.070 to 0.042 (96.0%; one-sided Wilcoxon *p* = 3.1 × 10^−5^) and reduces mean core-parameter error from 16.6% to 8.8% (*p* = 6.1 × 10^−5^). A sensitivity analysis on 72 viable specimens reveals that *τ*_*w*_ and *α* are the most consequential parameters for XOR twin accuracy, while *v*_scale_ and *R*_off_ are both hard to identify and tolerant to error. These results show that fungal digital twins can already narrow the search for viable computational substrates, partially recover the excitable dynamics that govern them, and support small-scale specimen-specific refinement without yet claiming full XOR transfer.

## Introduction

Fungal computing treats living mycelial networks as excitable, adaptive substrates for information processing (Adamatzky, 2018; Adamatzky et al., 2021a,b; Adamatzky, 2023). Mycelia generate structured electrical activity that has been compared to spiking patterns in neural systems (Adamatzky, 2022). They respond to stimulation and can realize logic operations through the propagation and interaction of signals in space and time (Adamatzky et al., 2020; Roberts and Adamatzky, 2022). They also exhibit memristive behavior, meaning their conductance depends on the history of applied voltage, making them attractive as low-energy, self-organizing computing materials (Beasley et al., 2021; LaRocco et al., 2025; Chua, 2003; Strukov et al., 2008). Recent work has modeled mycelium growth explicitly toward reservoir computing (Jaeger and Haas, 2004; Maass et al., 2002; Tompris et al., 2025), and morphologically tunable mycelium chips have been proposed as physical reservoir substrates (Telhan et al., 2025).

The central obstacle to reproducible fungal computing is variability. Different fungal specimens differ in network topology, excitability, recovery time scales, and adaptive conductance. As a result, an electrode configuration that produces useful logic on one specimen may not transfer to another. Overcoming this variability requires methods to both characterize individual specimens and predict which computational behaviors they can support.

This motivates *digital twins*: computational surrogates that can be probed and optimized *in silico* before physical intervention. More generally, digital twins are useful in biology when they support intervention planning while making clear which latent properties are identifiable from finite observations (Wu and Koelzer, 2024; Clancy and Landman, 2026). In that sense, fungal twin construction is also a problem of computer-model calibration and simulation-based inference (Kennedy and O’Hagan, 2001; Cranmer et al., 2020). A related approach in agent-based modeling uses machine learning surrogates to calibrate complex simulators from observed data (Lamperti et al., 2018). Our work differs from these general calibration frameworks in that the target is a living substrate whose computational function is emergent and task-dependent.

Several existing approaches address parts of this problem.

Reservoir computing with mycelium has been explored both in simulation (Tompris et al., 2025) and in hardware (Telhan et al., 2025), but these approaches typically treat the substrate as a fixed black box rather than attempting to infer its internal dynamics. Boolean logic gates in fungal colonies have been demonstrated experimentally (Adamatzky et al., 2020; Roberts and Adamatzky, 2022), but without a systematic framework for predicting which specimens will support which gates. Our work bridges these efforts by providing a computational pipeline that connects substrate characterization to logic-gate viability.

The specific logic operation we study is XOR (exclusive- or), chosen because it is linearly inseparable and therefore provides a nontrivial test of nonlinear computation. Unlike AND or OR gates, XOR requires that the substrate nonlinearly combines its inputs, making it a stringent probe of whether a fungal network can perform more than simple signal summation. Demonstrating that digital twins can predict XOR viability is therefore a meaningful step toward general-purpose fungal computing.

This paper asks five questions. First, which fungal parameter regimes are viable substrates for XOR computation? Second, do standardized electrical probes contain enough information to recover the latent parameters of the model? Third, which characterization protocols matter most for that recovery? Fourth, can ML predictions be improved by specimen-specific waveform matching on a reduced validation set? Fifth, which biophysical parameters most critically affect the accuracy of digital twins when they are used for XOR computation? The contribution is therefore a computational framework for substrate discovery and calibration to help inform future studies of full end-to-end XOR gate transfer.

## Methods

### Fungal Network Model

The fungal substrate is modeled as a random geometric graph *G* = (*V, E*) embedded in a 20 × 20 mm spatial domain. Nodes represent hyphal junctions and are placed uniformly at random; edges connect nodes within a coupling radius of 5 mm (normalized to 0.25 in unit-square coordinates). Each node *i* follows FitzHugh–Nagumo excitable dynamics (FitzHugh, 1961; Nagumo et al., 1962; Izhikevich and FitzHugh, 2006):

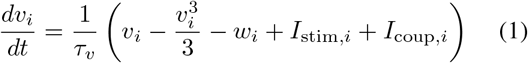

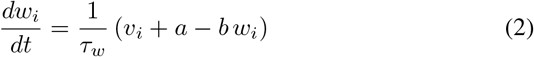

where *v*_*i*_ is the fast voltage variable, *w*_*i*_ is the slow recovery variable, *I*_stim,*i*_ is the external stimulation current, and *I*_coup,*i*_ is the coupling current from neighboring nodes. The parameters *τ*_*v*_ and *τ*_*w*_ set the fast and slow time scales; *a* and *b* control excitability and recovery.

Each edge (*i, j*) ∈ *E* carries a memristive state *m*_*ij*_ ∈ [0, 1] that modulates the effective conductance:

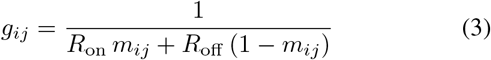

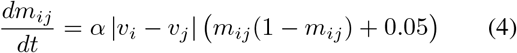

where *R*_on_ and *R*_off_ are the low- and high-resistance states, α is the memristive adaptation rate, and the small constant prevents the state from stalling at boundaries. The coupling current for node *i* is then *I*_coup,*i*_ = Σ_*j*_ ∈ 𝒩_(*i*)_ *g*_*ij*_(*v*_*j*_ − *v*_*i*_). Electrode stimulation enters each node via a Lorentzian spatial decay: *I*_stim,*i*_ = *V*_elec_*/*(1 + (*d*_*i*_*/r*)^2^), where *d*_*i*_ is the distance from node *i* to the electrode and *r* is the coupling radius.

The model has eight latent biophysical parameters: *τ*_*v*_, *τ*_*w*_, *a, b, v*_scale_, *R*_off_, *R*_on_, and *α*. The parameter *v*_scale_ is a global voltage scaling factor. Together these parameters capture the essential physics of fungal excitable networks: the FHN parameters (*τ*_*v*_, *τ*_*w*_, *a, b*) govern the threshold, shape, and recovery of action potential-like pulses, while the memristive parameters (*R*_off_, *R*_on_, *α*) control history-dependent conductance changes akin to those observed in fungal tissue and in the broader memristor literature (Beasley et al., 2021; Chua, 2003; Strukov et al., 2008). Network size varied from 20 to 120 nodes across experiments, yielding graphs with mean degree proportional to the node density at fixed coupling radius. Simulations were run with adaptive ODE integration in Python, with a safety mechanism that halts integration when state variables exceed physical bounds.

### Viable XOR Range Discovery

To identify parameter regimes that support logic, we ran a systematic optimization study on 160 simulated specimens spanning eight network sizes (20, 30, 40, 50, 60, 80, 100, 120 nodes; 20 trials per size). For each specimen, Bayesian optimization with a Gaussian-process surrogate (Snoek et al., 2012; Shahriari et al., 2015) searched over electrode positions (two input electrodes, one output probe), stimulus voltage, pulse duration, and inter-input delay to maximize XOR performance. XOR accuracy was scored by applying all four input combinations (00, 01, 10, 11) and measuring whether the output voltage at the probe exceeded a threshold for the two “on” inputs and fell below it for the two “off” inputs.

A second tuning stage optimized the eight biophysical parameters while holding the optimized gate geometry fixed. This two-stage design separates the question of “where to stimulate” from “what physics to use,” ensuring that the viable range analysis captures the intrinsic substrate properties rather than electrode-specific artifacts.

XOR performance was quantified by a composite score that combines classification accuracy across the four input combinations with the margin between “on” and “off” output voltages. A higher score indicates both correct classification and a wider voltage gap, which corresponds to a more robust gate that is less sensitive to noise. Viability was defined as membership in the top quartile of tuned XOR scores (threshold *≥* 0.0627), yielding 40 viable specimens that define a task-specific prior over fungal parameters likely to support XOR-like behavior.

### Characterization Dataset

To build an inverse model, we generated 400 simulated specimens (50 per network size across the same eight graph sizes). Each specimen was probed with three characterization protocols (Figure 1):

**Figure 1:**
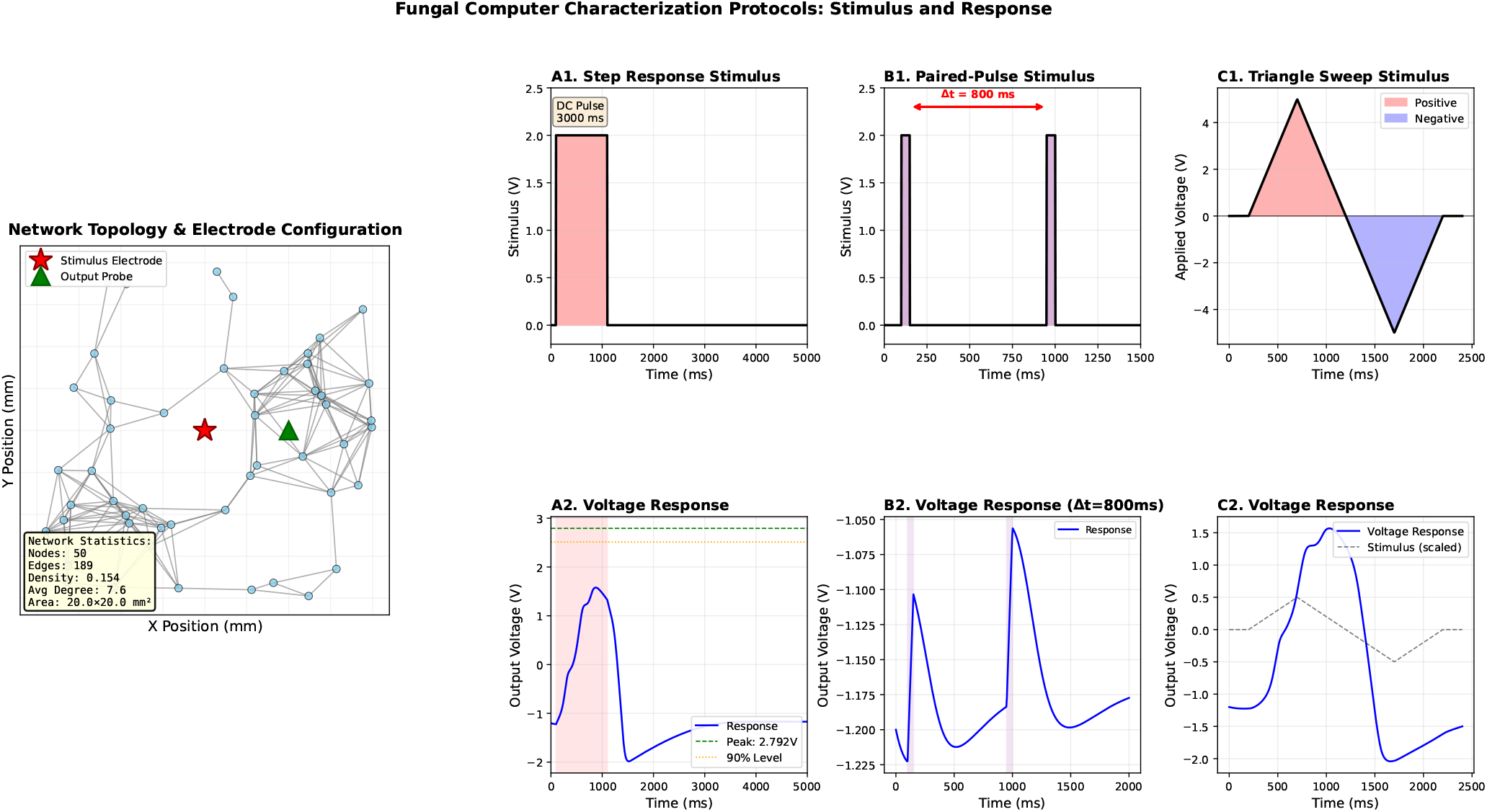
Characterization protocols used to probe the fungal digital twin model. The left panel shows a representative network and stimulation geometry; the right panels illustrate the step-response, paired-pulse, and triangle-sweep protocols and the waveform statistics extracted from each response.

#### Step response

A sustained voltage pulse (2 V, 3000 ms duration) applied through a single electrode, with the response recorded at a probe 5 mm away. This protocol measures the network’s transient excitation and recovery dynaics. Thirteen features were extracted, including peak amplitude, time to peak, half-decay time, activation speed, latency, and asymmetry index.

#### Paired-pulse

Two identical short pulses (2 V, 50 ms each) separated by interpulse intervals of 200, 800, and 2000 ms. This protocol probes recovery-dependent facilitation or depression. Eighteen features were extracted per delay (54 total plus 4 aggregate features), including peak ratios, waveform correlation, and facilitation indices.

#### Triangle sweep

A linear voltage ramp up to 5 V and back at a rate of 0.01 V/ms, probing the current–voltage hysteresis loop characteristic of memristive behavior. Twenty-three features were extracted, including hysteresis area, rectification index, loop eccentricity, and smoothness index.

In total, 94 features describe each specimen. The characterization study intentionally sampled the full model parameter box rather than only the XOR-viable subset, so the resulting predictor is a broad identifiability model rather than a viable-range-specific one.

### Parameter Prediction and Ablation

We trained separate regressors for each target parameter using the 94-dimensional feature vectors. Two model families were compared: random forests (200 trees, max depth 20) and multilayer perceptrons (three hidden layers: 128, 64, 32 units). Performance was measured on a held-out 20% test split using *R*^2^ and RMSE. To identify which probes mattered most, we retrained the random forest models on seven feature subsets: step only (13 features), paired-pulse only (58), triangle only (23), and all pairwise and three-way protocol combinations.

### Sensitivity Analysis

To assess which parameters most critically affect twin-based XOR accuracy, we ran a perturbation study on 72 viable specimens (top 50% by baseline twin XOR accuracy). For each specimen, the true tuned parameters were perturbed one at a time at *±*5%, *±*10%, *±*20%, and *±*30% magnitudes, and the XOR accuracy of the resulting perturbed twin was recorded. The sensitivity metric is the drop in twin XOR accuracy relative to the unperturbed baseline, averaged over both directions of perturbation at each magnitude.

### Rediscovery Validation

To test whether inverse predictions are useful starting points for specimen-specific calibration, we ran a preliminary rediscovery study on the top-quartile optimized specimens with at most 50 nodes. Twins copied the exact network structure of each specimen, so the rediscovery task isolated the eight latent biophysical parameters rather than topology. Three conditions were evaluated: *oracle* (true tuned parameters), *ML only* (random-forest predictions from characterization features), and *ML+refine* (the same predictions followed by waveform-matching refinement). Refinement used dual annealing with local bounds centered on the ML prediction, with each bound spanning *±* 20% of the full parameter range. The objective minimized mismatch between specimen and twin waveforms collected under the same step, paired-pulse, and triangle-sweep protocols. One specimen produced non-finite characterization features and was excluded, leaving *n* = 15 specimens.

### Statistical Analysis

Held-out prediction quality was summarized with *R*^2^ and RMSE. Rediscovery comparisons between ML only and ML+refine used one-sided paired Wilcoxon signed-rank tests because the sample was small and the directional hypothesis was pre-specified: refinement should reduce waveform mismatch and parameter error. For rediscovery we report mean relative error across all eight parameters and across a core identifiable subset (*τ*_*v*_, *τ*_*w*_, *a, b, α*). Sensitivity analysis used mean accuracy drop averaged across both perturbation directions at each magnitude.

## Results

### XOR Viability Occupies a Constrained Region of Parameter Space

The systematic optimization study completed 160/160 runs successfully. Defining viability as the top quartile of tuned XOR scores yielded 40 viable specimens with threshold 0.0627. Viable specimens appeared across the full range of graph sizes, although the highest-scoring subset was somewhat enriched in larger networks (100–120 nodes), suggesting that larger graphs increase the probability of useful specimens without defining a single privileged topology.

Figure 2 summarizes the viable region. Compared to the full search bounds, the viable subset concentrates around *τ*_*v*_ *≈* 54 ms (full range 30–150), *τ*_*w*_ *≈* 894 ms (300–1600), *a ≈* 0.67 (0.5–0.8), *b ≈* 0.87 (0.7–1.0), *R*_off_ *≈* 182 Ω (50–300), *R*_on_ *≈*17.9 Ω (2–50), and α*≈* 0.009 (0.0001– 0.02). The viable range covers only 56% of the *τ*_*v*_ search range and 60% of *τ*_*w*_, indicating genuine constraint on the excitable time scales required for XOR. Viable stimulation geometry was also structured: input electrodes were separated by 7.8 *±* 6.4 mm, the mean input-to-output distance was 13.1 *±* 3.5 mm, mean voltage was 2.71 *±* 0.91 V, and mean pulse duration was 2869 *±* 1634 ms.

**Figure 2:**
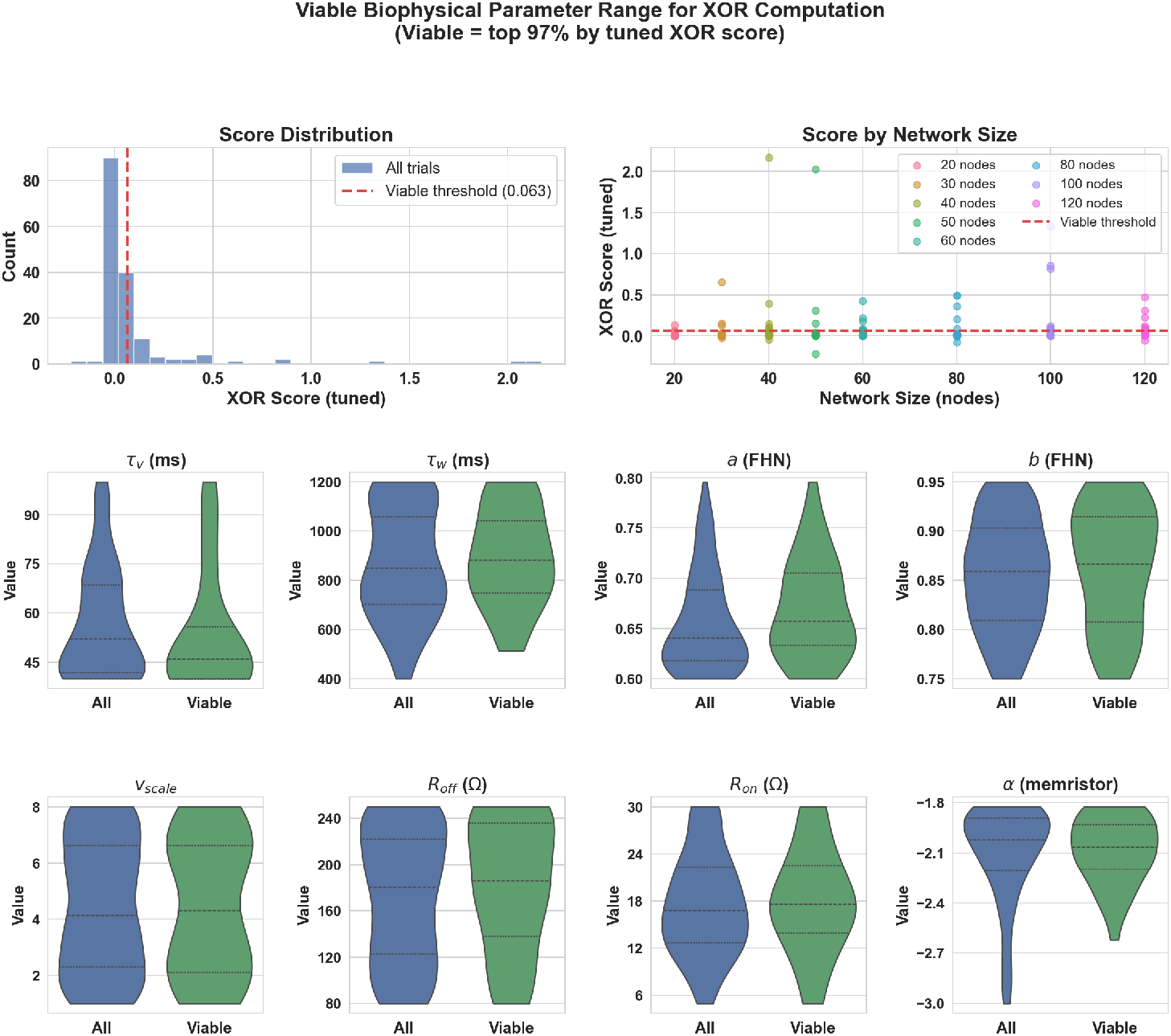
Distribution of tuned XOR scores and tuned parameter values for all optimized specimens versus the top quartile of viable XOR substrates. Viable specimens occupy a narrower parameter region than the full search space, providing a computational prior for subsequent identification studies.

Four of the eight parameter marginals showed bimodality (bimodality coefficient *>* 0.555): *τ*_*v*_, *b, v*_scale_, and *R*_off_. This multimodality suggests that XOR-supporting fungi are better interpreted as a family of dynamical regimes than as a single optimum, consistent with the view that different combinations of excitability and plasticity can produce equivalent computational affordances. For an experimental program, this means that characterizing a specimen into one of these dynamical modes may be more informative than trying to match a single target parameter vector.

In terms of the viability rate, 25% of specimens (by construction, as the top quartile) were classified as viable, but the absolute XOR score distribution was heavily right-skewed: the best specimen scored 65 × higher than the viability threshold. This suggests that while many parameter combinations can produce marginal XOR behavior, truly robust gates are rare and require precise alignment of excitable dynamics and electrode geometry.

### Electrical Probes Recover Some Parameters Reliably but Not All

All 400 characterization simulations completed successfully. Random forests clearly outperformed multilayer perceptrons and were therefore used in all downstream analyses.

The strongest global predictions were for the two time constants and the excitability threshold: *R*^2^ = 0.912 for *τ*_*v*_, 0.816 for *τ*_*w*_, and 0.717 for *a* (Table 1). Parameter *b* was only moderately predictable (*R*^2^ = 0.342), while *α* remained weak but above chance (*R*^2^ = 0.396). In contrast, *v*_scale_, *R*_on_, and *R*_off_ were not recovered reliably from the present protocols (*R*^2^*≤* 0.02). This is an important negative result: the current stimulation set supports partial, not complete, identifiability.

**Table 1:** Held-out random forest performance overall and within each parameter’s viable interval. Viable intervals are taken from the XOR-viable range analysis. Lower viable-range *R*^2^ values should be interpreted carefully because the target variance is smaller inside a restricted interval.

| Parameter | Overall $R^2$ | Overall RMSE | Viable $n$ | Viable $R^2$ | Viable RMSE |
| --- | --- | --- | --- | --- | --- |
| $\tau_v$ | 0.912 | 10.566 | 37 | 0.746 | 8.857 |
| $\tau_w$ | 0.816 | 145.757 | 58 | 0.730 | 127.935 |
| $a$ | 0.717 | 0.044 | 47 | 0.065 | 0.047 |
| $b$ | 0.342 | 0.068 | 63 | 0.210 | 0.056 |
| $v_{scale}$ | -0.007 | 2.501 | 71 | -0.031 | 2.218 |
| $R_{off}$ | -0.244 | 81.195 | 62 | -0.196 | 63.808 |
| $R_{on}$ | 0.017 | 13.538 | 39 | -1.340 | 12.259 |
| $\alpha$ | 0.396 | 0.003 | 30 | -0.038 | 0.004 |

Breaking performance out by viable interval clarifies how these predictors behave where XOR is actually relevant. Absolute RMSE within the viable interval is similar to or lower than the overall RMSE for six of the eight parameters, including *τ*_*v*_, *τ*_*w*_, *b, v*_scale_, *R*_off_, and *R*_on_. However, viable-range *R*^2^ is often lower than overall *R*^2^ because the viable subset occupies a narrower slice of parameter space. In practice, this means the predictors can still be locally useful inside the XOR-relevant regime even when variance-explained scores deteriorate under range restriction.

### Different Protocols Capture Different Latent Dynamics

Protocol ablation reveals a clear division of labor across the stimulation set (Figure 3). The full 94-feature model gave the best overall mean performance, but no single protocol dominated every target. Step-response features carried most of the information for *τ*_*v*_ (*R*^2^ = 0.825 step-only vs. 0.865 all). Paired-pulse features were most informative for *a* (*R*^2^ = 0.692 paired-pulse-only), indicating that short-timescale recovery and facilitation are especially diagnostic of the excitation threshold. The combination of step response and triangle sweep was best for *τ*_*w*_ (*R*^2^ = 0.752), *α* (*R*^2^ = 0.374), and *R*_off_, suggesting that slower recovery and memristive properties are better exposed by combining transient and hysteretic probes. Parameters *v*_scale_ and the resistance pair remained difficult across all ablations, reinforcing the interpretation that the present protocol library underconstrains those aspects of the model.

**Figure 3:**
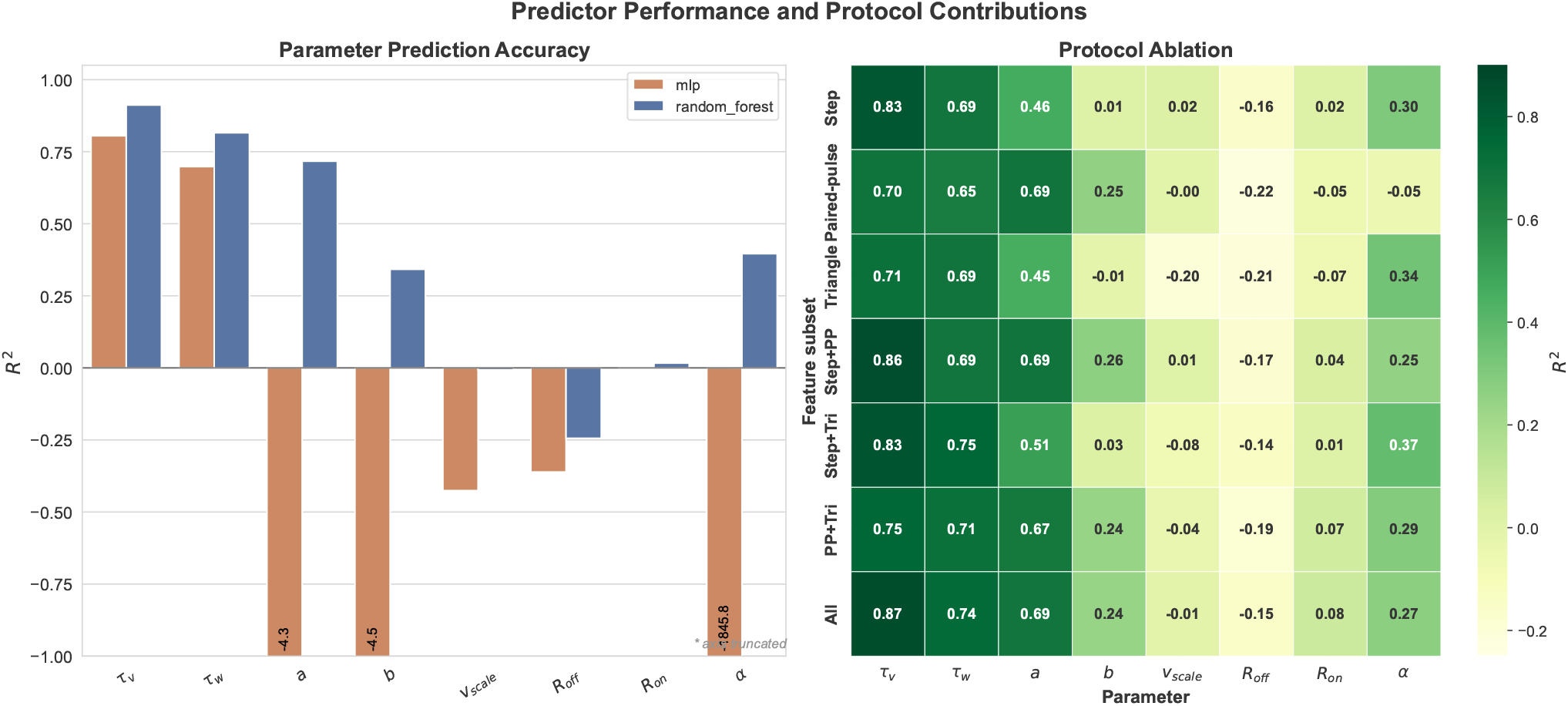
Left: held-out *R*^2^ for random forest and MLP predictors across the eight biophysical parameters. Right: protocol-ablation heatmap showing how prediction quality depends on which characterization protocols are included.

An important practical implication is that the paired-pulse protocol alone achieves *R*^2^ = 0.692 for *a* while step response alone reaches only 0.461. This suggests that if experimenters must choose a single additional protocol beyond step response, the paired-pulse protocol offers the largest marginal gain for excitability-related parameters.

### Sensitivity Analysis Reveals a Hierarchy of Parameter Importance

The sensitivity analysis on 72 viable specimens reveals a clear hierarchy of parameter importance for XOR computation (Figure 4). At *±* 30% perturbation, the recovery time constant *τ*_*w*_ caused the largest mean accuracy drop (0.021), followed by *α* (0.014), *b* (0.014), *a* (0.012), and *τ*_*v*_ (0.012). In contrast, *R*_off_ and *v*_scale_ had negligible or negative impact, meaning that perturbing them by *±* 30% barely affected XOR accuracy.

**Figure 4:**
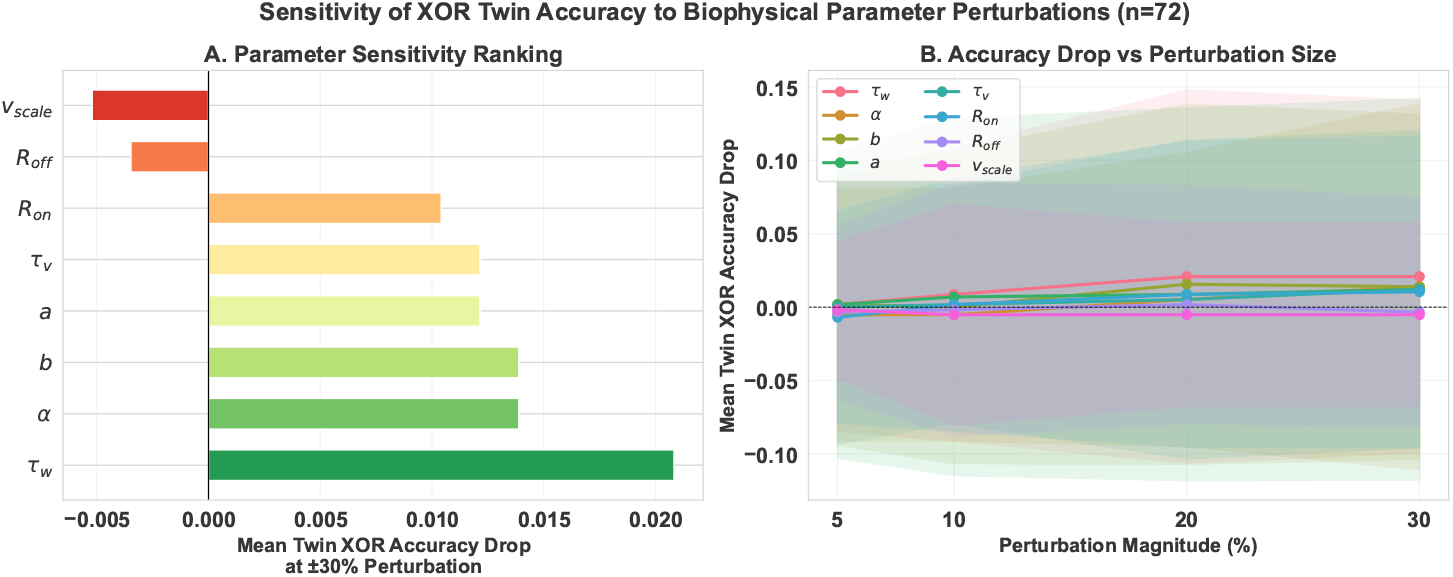
Sensitivity of twin XOR accuracy to single-parameter perturbations across 72 viable specimens. Left: mean accuracy drop at *±* 30% perturbation (tornado chart). Right: accuracy drop versus perturbation magnitude for all eight parameters. The recovery time constant *τ*_*w*_ and memristive adaptation rate *α* are the most consequential for XOR accuracy.

This result connects identifiability and functional relevance. The parameters that are easiest to predict from characterization data (*τ*_*v*_, *τ*_*w*_, *a*) are also among the most consequential for XOR accuracy. Conversely, *v*_scale_ and *R*_off_, which are poorly identifiable, are also the least consequential for XOR. This is a reassuring alignment: the workflow has the highest fidelity on the parameters that matter most for the target computation, and the parameters it recovers poorly happen to be those where errors are tolerable.

The one exception to this pattern is *α*: it is moderately difficult to predict from characterization data (*R*^2^ = 0.396) but ranks as the second most consequential parameter for XOR accuracy. This identifies memristive adaptation rate as a priority target for future protocol design—a probe specifically sensitive to *α* would yield the largest improvement in twin utility for logic applications.

The sensitivity hierarchy also has implications for how we interpret the viable range. The viable region for *τ*_*w*_ spans 60% of the full search range, but *τ*_*w*_ perturbations cause the largest XOR accuracy drops. This means that *τ*_*w*_ must be both correctly chosen (to fall in the viable range) and precisely estimated (because errors propagate to twin accuracy), making it the single most important parameter in the entire digital-twin pipeline. In contrast, *v*_scale_ has a viable range spanning 86% of the full search bounds and minimal sensitivity to perturbation, suggesting it plays a permissive rather than instructive role in XOR computation.

### Waveform-Guided Refinement Improves Rediscovery

Rediscovery used 16 optimized specimens with 20–50 nodes; one specimen was excluded after producing non-finite characterization features, leaving 15 paired cases. Table 2 summarizes the results. Oracle twins recovered zero error by construction. ML-only initialization yielded mean waveform mismatch 1.070 *±* 1.625 and mean core-parameter error 16.6 *±* 5.9%. After waveform-matching refinement, mismatch fell to 0.042 *±* 0.071 and core-parameter error fell to 8.8 *±* 4.4%. Across all eight parameters, mean relative error dropped from 31.6 *±* 18.2% to 23.8 *±* 20.3%.

**Table 2:** Rediscovery validation on 15 optimized specimens after excluding one numerical failure.

| Condition | $n$ | Waveform mismatch | Mean rel. err. (all) | Mean rel. err. (core) |
| --- | --- | --- | --- | --- |
| Oracle | 15 | $0.000 \pm 0.000$ | $0.0 \pm 0.0$ | $0.0 \pm 0.0$ |
| ML only | 15 | $1.070 \pm 1.625$ | $31.6 \pm 18.2$ | $16.6 \pm 5.9$ |
| ML+refine | 15 | $0.042 \pm 0.071$ | $23.8 \pm 20.3$ | $8.8 \pm 4.4$ |

These reductions were statistically strong under one-sided paired Wilcoxon tests. ML+refine outperformed ML only for waveform mismatch (*W* = 120, *p* = 3.1 × 10^−5^), mean relative error across all parameters (*W* = 118, *p* = 9.2 × 10^−5^), and mean relative error across the core identifiable subset (*W* = 119, *p* = 6.1 × 10^−5^). The average waveform mismatch reduction was 96.0%, while mean core-parameter error fell by 46.9%.

At the parameter level, refinement substantially improved *τ*_*w*_ (10.98%*→* 3.20%), *a* (7.04% *→*3.08%), *R*_on_ (54.83%*→* 14.87%), and *α* (49.90%*→* 25.33%) with one-sided Wilcoxon *p <* 2.2 × 10^−4^ in each case. Improvement for *τ*_*v*_ was smaller and not significant at this sample size, while *b* and *R*_off_ were essentially unchanged. Parameter *v*_scale_ worsened after refinement (78.58%*→* 94.81%), which is consistent with the earlier predictor analysis showing that it is weakly constrained by the present protocols. Taken together with the sensitivity analysis, this worsening is less alarming than it might appear: *v*_scale_ is the parameter whose perturbation has the least impact on XOR accuracy.

## Discussion

### Summary of Contributions

The main result of this paper is that fungal digital twins are already useful at three levels: they can discover viable computational regimes, they can partially infer the excitable and adaptive dynamics that make those regimes possible, and they can support small-scale specimen-specific refinement by waveform matching. The optimization study narrows the search from the full fungal-like parameter box to a viable XOR-supporting subspace occupying roughly half the original ranges of the key time-scale parameters. The inverse-model study shows that standardized electrical probes recover some latent parameters reliably, especially the time constants and the main excitation threshold. The preliminary rediscovery study then shows that those predictions are good enough to initialize a second calibration stage that substantially improves twin fidelity. The sensitivity analysis closes the loop by confirming that the parameters the work-flow recovers best are the ones that matter most for XOR computation.

### Fungal Computing as an Artificial-Life Problem

This supports a specific view of fungal computing as an artificial-life problem. The fungal substrate is an adaptive physical medium whose computational affordances emerge from the interaction of topology, excitability, and plasticity. The digital twin is therefore valuable because it narrows the design space to regions where useful behavior is likely and identifies which hidden variables can be recovered from experimentally accessible probes.

The multimodality observed in the viable parameter space is particularly relevant. Four of eight parameters showed bimodal distributions among XOR-viable specimens, suggesting that there are multiple dynamical regimes that can produce equivalent XOR behavior. This resonates with a central theme: that functional equivalence across structurally different substrates can arise from the interaction of multiple adaptive components. Understanding these equivalence classes may be more valuable for fungal computing than finding a single “optimal” parameter set, because it broadens the pool of natural specimens that could serve as viable substrates.

### Identifiability, Sensitivity, and Protocol Design

The negative results are equally important. Three parameters (*v*_scale_, *R*_on_, *R*_off_) are poorly constrained by the present protocol suite at the prediction stage, and only one of them (*R*_on_) improves reliably under waveform refinement. This indicates a structural mismatch between what the model treats as important and what the current probes actually observe. This may be seen as a weakness of our approach; but can also be framed as a contribution. The ablation study shows that protocol design and identifiability are coupled, and the rediscovery study shows which parameters remain underdetermined even after specimen-specific calibration.

The sensitivity analysis adds a crucial pragmatic dimension: it reveals that the parameters hardest to identify (*v*_scale_, *R*_off_) are precisely the ones least consequential for XOR performance. Parameters that have little effect on the output waveform are intrinsically difficult to recover from that waveform, but they also matter less for downstream computation. The practical consequence is that the current protocol suite, despite its identifiability gaps, is already well-matched to the requirements of XOR twin construction. The one parameter that breaks this convenient alignment is *α*, which is moderately hard to predict but functionally important. This identifies memristive adaptation rate as the single highest-priority target for future protocol design.

The protocol ablation provides concrete guidance for experimentalists: the paired-pulse protocol is the most informative single addition to a step-response baseline, particularly for recovering the excitation threshold *a*. If a third protocol is feasible, the triangle sweep adds the most information about *τ*_*w*_ and *α* when combined with step response. This suggests a practical ordering for experimental deployment: step response first, paired-pulse second, triangle sweep third.

### Limitations and Scope

There are clear limits to the present claim set. First, the characterization dataset samples the full parameter box rather than only the XOR-viable subset. That choice is useful for establishing baseline identifiability across the whole model family, but viable-range-conditioned retraining is the natural next step for task-specific twin construction. Second, the rediscovery study keeps network topology fixed by copying the specimen graph into the twin. The current results therefore isolate parameter calibration rather than full structure-and-parameter rediscovery. Recovering network topology from electrical measurements is a substantially harder inverse problem that remains open. Third, this paper does not test whether calibrated twins reliably transfer optimized XOR gates back to unknown specimens. That stronger end-to-end claim requires demonstrating that a gate designed *in silico* on the twin actually works when applied to the physical specimen, and remains a journal-scale extension. Fourth, the sensitivity study uses a one-at-a-time perturbation design, which does not capture interaction effects between parameters. Higher-order sensitivity methods such as Sobol indices (Sobol, 2001) would provide a more complete picture but require significantly more computational budget.

### Future Directions

The limitations above define the next experiments. The viable-range analysis suggests that both *R*^2^ and RMSE should be reported when moving to local calibration problems, because range restriction can depress *R*^2^ even when absolute error improves. The protocol ablation and rediscovery results point to richer conductance-sensitive probes— such as impedance spectroscopy or frequency-dependent stimulation—as the fastest route to resolving *v*_scale_ and *R*_off_, and especially *α*.

The hybrid ML-plus-refinement workflow used here aligns naturally with broader simulation-based inference strategies (Cranmer et al., 2020). An amortized posterior estimator, for example a neural density estimator trained on the characterization dataset, could replace the point-prediction random forest with a full posterior over parameters and uncertainty-aware calibration (Papamakarios et al., 2021). Similarly, Bayesian optimization of the characterization protocol itself—choosing which probes to run next based on which parameters remain uncertain—could substantially reduce the number of experiments needed per specimen (Snoek et al., 2012; Shahriari et al., 2015).

A critical next step is experimental validation on physical fungal specimens. The characterization protocols described here (step response, paired-pulse, triangle sweep) are standard electrophysiological procedures that can be applied to real mycelial networks with minimal modification. The key empirical question is whether the FHN-memristor model, despite its simplifications, captures enough of the real substrate dynamics to produce useful digital twins. Even partial success—recovering the correct time scales and excitability regime, if not the precise resistance values—would enable the twin-guided optimization workflow to propose electrode configurations that are at least approximately correct for a given specimen.

Beyond XOR, the framework generalizes to other logic operations and to reservoir computing (Maass et al., 2002; Jaeger and Haas, 2004), where the goal shifts from gate fabrication to maximizing the computational capacity of the substrate. The characterization protocols and inverse models developed here could be adapted to predict reservoir properties such as kernel rank and generalization capability, potentially enabling digital-twin-guided reservoir design.

Together, these results make a complete contribution: viable-regime discovery, protocol-based identifiability analysis, a sensitivity hierarchy linking identifiability to functional importance, and a reduced but statistically supported rediscovery validation.

## Conclusion

We have presented a digital-twin workflow for fungal excitable networks to enable reproducible fungal computing. The framework is designed to generalize beyond XOR to other logic gates. The characterization protocols, inverse models, and sensitivity methods apply to any computational task that can be scored from the output of a fungal network model. As the community moves toward more complex operations and larger-scale reservoir computing, the digital-twin approach described here provides the infrastructure for systematic substrate selection, characterization, and calibration.

## Notes

### Competing Interest Statement

The authors have declared no competing interest.

